# Surface detection of SARS-CoV-2 by lateral flow LAMP

**DOI:** 10.1101/2022.04.04.487067

**Authors:** Isabelle Dahl Acker, Mark Joseph Ware, John Russell Bracht

**Affiliations:** Department of Health Studies, American University, Washington DC 20016; Biology Department, American University, Washington DC 20016

## Abstract

Slowing the transmission of SARS-CoV-2 requires rapid and accurate diagnostic testing. Toward this end, loop-mediated isothermal amplification (LAMP), an isothermal genomic detection method, offers great promise but the readout tends to be difficult because it does not generate linear DNA products. Rapid antigen tests are coupled to lateral flow strips, with one (negative) or two (positive) bands providing simple rapid readout, but are less sensitive than genomic amplification methods. To address the need for a genomic amplification method that can be visualized on a lateral flow strip, we developed a novel strand-displacement probe. In this work we validate this pipeline for purified RNA, intact virus, and even virus deposited onto a surface. We demonstrate robust sensitivity (100 genomic copies) and and we demonstrate the utility of our assay as a surveillance system, with the capability to detect viral particles from surfaces, even after a week of complete dry-down. Our innovation couples the diagnostic advantages of a nucleic acid amplification test (NAAT) with the simplicity of lateral-flow readouts.

## Introduction

The worldwide COVID-19 pandemic, driven by SARS-CoV-2 virus, is a particularly potent challenge to healthcare infrastructure, posing enhanced risk for individuals with preexisting risk factors such as cardiovascular disease, cancer, and obesity [1–4]. Even with the advent of highly efficacious vaccines [5,6], the spread of SARS-CoV-2 continues due to the emergence of new variants and vaccine hesitancy [7]. Thus diagnostic and surveillance testing are ongoing needs for the foreseeable future. Currently, the main strategy is to use either PCR-based methods [8] or rapid tests, which involve viral antigen detection with antibodies [9]. PCR methods are highly accurate because they are based on nucleic acid amplification; however, the assay must be run in a lab, with expensive, specialized equipment and well-trained lab technicians. By contrast, antigen tests are simpler to run, and yield faster results, but are less sensitive than PCR [9]. Isothermal amplification methods, which are related to PCR but are generally faster and can be run without complex equipment [10], offer the potential to bridge the best of both worlds. However, these isothermal methods often lack a simple readout equivalent to an antigen test.

We decided to use the loop-mediated isothermal amplification (LAMP) method. LAMP amplifies nucleic acid with primers and DNA polymerase-mediated strand displacement [11]. Unlike PCR, LAMP amplifies at a constant temperature and generates significant amplification in 10-15 minutes [12]. Unlike PCR, the polymerase (Bst) is not generally inhibited by compounds found in the clinical samples so RNA purification is often not required [13] making the setup simpler. However, readout methods for LAMP have been cumbersome due to the unorthodox zigzag structure of the DNA produced, precluding traditional gel electrophoresis [11]. Colorimetric readouts have been developed that rely on a sample’s pH changes to indicate amplified product [14]. However, this method is prone to both false positive and false negative errors because some samples display high or low pH independently of LAMP amplification, or may exhibit buffering capacity preventing pH change in spite of successful amplification [15].

A detection method directly linked to amplified DNA would avoid these errors. The Ellington laboratory recently described the strand-displacement probe [16], a double-stranded oligo that can directly link LAMP-amplified DNA to a lateral flow strip through displacement hybridization [17]. The double-stranded oligo probe sequence is constructed of a ‘reporter’ strand with FAM (6-carboxyfluorescein) at its 3’ end and a ‘quencher’ oligo with a quencher moiety at its 5’ end. Thus when hybridized the FAM fluorescent signal is quenched. The reporter has an 11-bp 5’ ‘toehold’ which is left single-stranded (Table 1). In the presence of the LAMP reaction product with a loop complementary to this toehold (and overall, to the reporter strand), a process of strand-displacement occurs which both attaches FAM to the LAMP product and separates FAM from its quencher (Fig. 1). The resultant product is suitable for detection both by fluorescent readout and on a lateral flow strip.

**Table 1.**
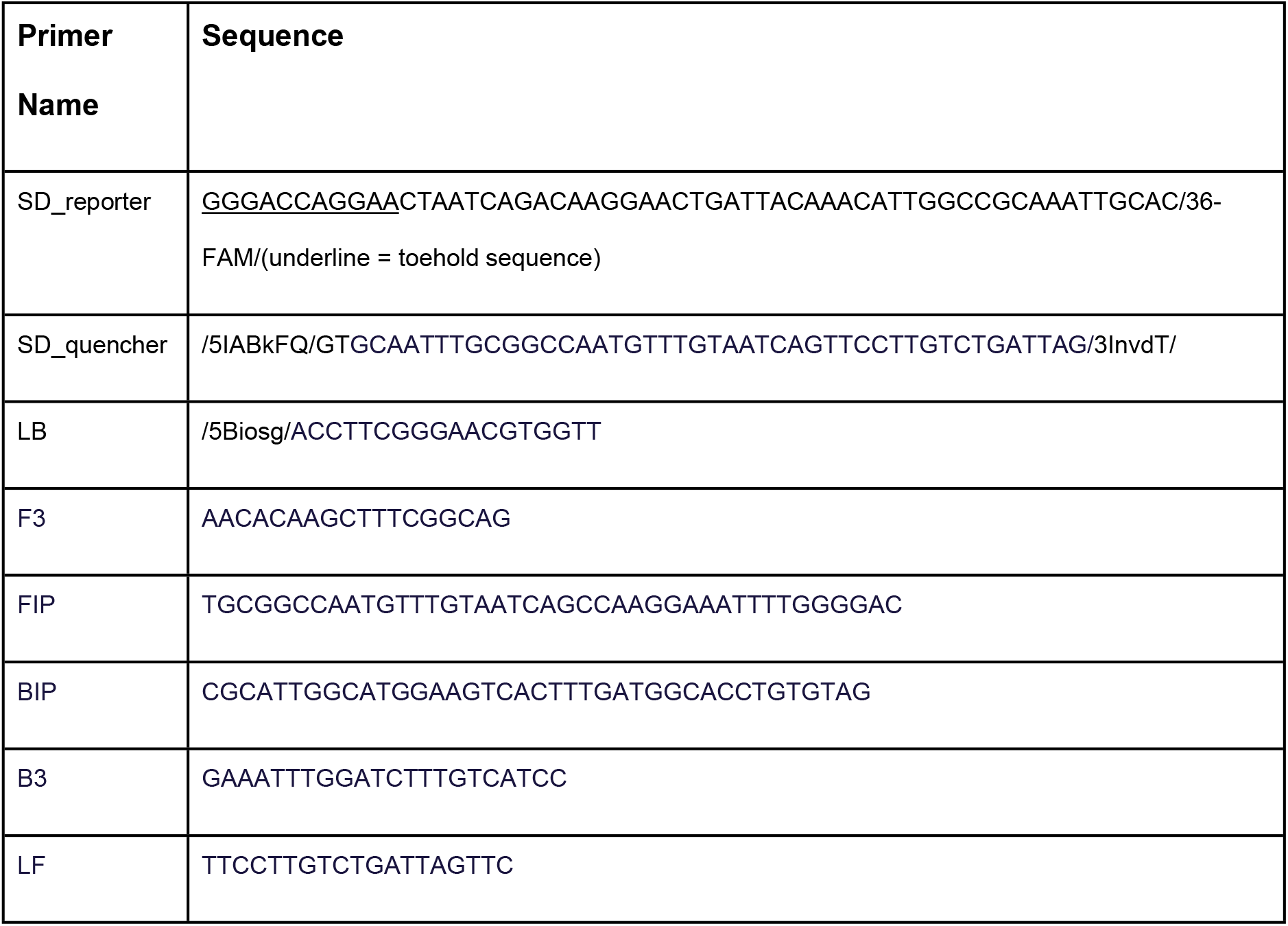
Primer sequences used in this study.

**Fig. 1.**
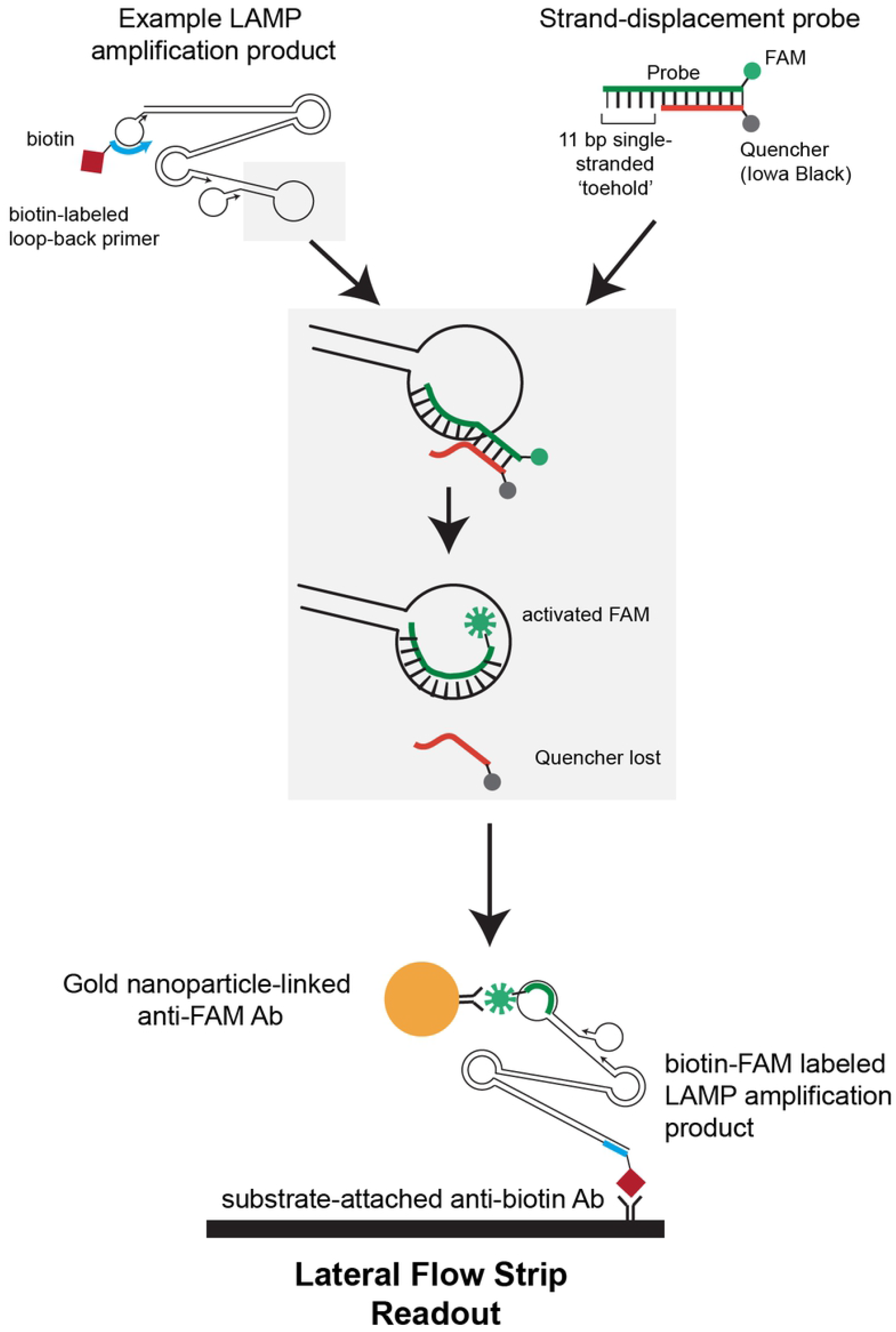
Overview of strand-displacement probe function. The strand-displacement probe has an 11-bp single-stranded toehold complementary to a loop of the LAMP amplification product. In the presence of LAMP products, thermodynamic-driven strand displacement leads to displacement of the quencher-strand, and activation of FAM fluorescence. Because the loop-back LAMP primer includes biotin, the resulting FAM-biotin linkage can be detected on a lateral flow assay or by measuring activated FAM fluorescence.

A commonly used lateral flow strip design detects products with both biotin and FAM to immobilize a gold nanoparticle to the test band [16]. One LAMP primer (in our case, LB) is linked to biotin while as described the FAM is attached to the strand-displacement probe’s reporter strand. Here we demonstrate that this approach successfully detects LAMP amplification products as simple-to-read bands while making minimal impact on reaction setup complexity. We demonstrate that the system exhibits excellent sensitivity and specificity, and that it can detect raw virus without nucleic acid extraction; furthermore, we show that it readily detects the virus dried onto surfaces. Thus, our results pave the way for point-of-care applications or on-site rapid surface surveillance.

## Methods

### LAMP

All LAMP reactions use the following recipe:

12.5 μL 2X LAMP mix (New England Biolabs Cat # E1700S or M1800S for colorimetric readout)

2.5 μL 10X primer mix

2.5 μL 1μM annealed probe*

7.5 μL template plus water

Incubate 60°C for 30 min, then readout color, if using dipstick (Milenia HybriDetect MGHD 1) add 10 μL directly to pad and immerse in 200 μL PBST. Incubate 2-5 min for band formation.

*to make annealed probe: primers mixed at equimolar 100 μM concentration, heated to 95°C for 3 min in a heatblock, then allowed to cool to room temperature when the block was switched off; this took approximately 20 min.

### Method for Fig. 2

**Fig. 2.**
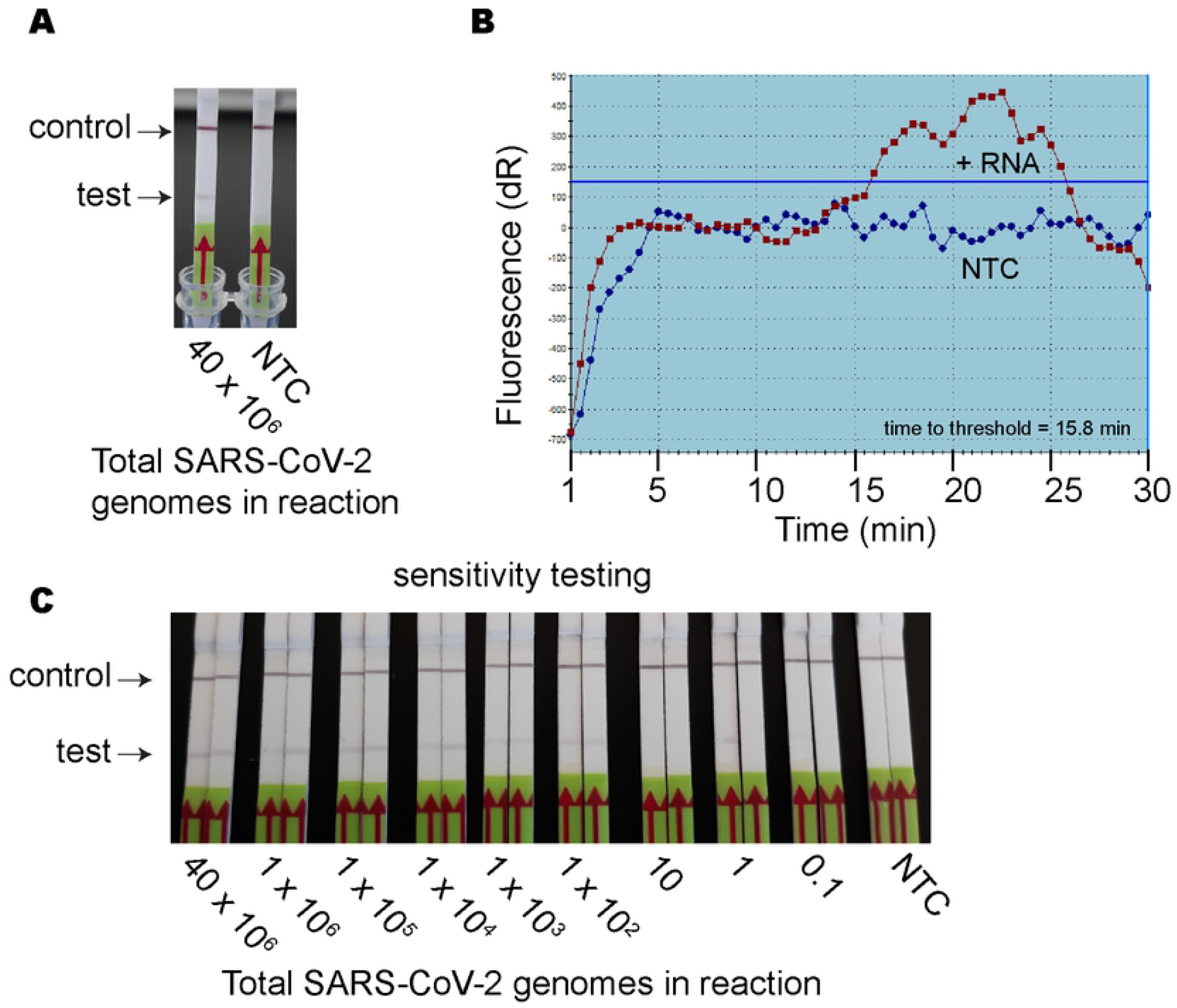
Proof of principle using lateral flow and fluorescence readouts, and sensitivity testing. A, Lateral flow readout of base SARS-CoV-2 genome concentration (40*10^6^); this concentration was used for all other experiments. No template control (NTC) is included on the right. B, Fluorescence readout of FAM (y-axis) across time (x-axis). Rise in RNA (positive control) readout is due to the displacement of the probe’s quencher while the subsequent drop is attributed to the pH drop generated by extensive DNA synthesis: FAM is very pH sensitive and quenches at lower pH. C, Serial dilutions of decreasing amounts (in viral molecules) of inactivated SARS-CoV-2 virus on lateral flow assay readouts. NTC shows no template control, each assay shows control band (top) and detection band (bottom), and each dilution was tested in duplicate.

Gamma-inactivated SARS-CoV-2 was obtained from BEI Resources, NIAID, (https://www.beiresources.org/), catalog # NR-52287. This material is supplied at 1.75 × 10^9^ genome equivalents per mL. We extracted RNA from 300μL of this gamma-inactivated virus with Zymo DNA/RNA Quick Miniprep Plus kit (Cat #D7003) to yield a stock of 5.25 ×10^6^ genomes/μL. Every LAMP reaction uses 7.5μL of RNA plus water, so without dilution this stock material yields 40 × 10^6^ genomes in a reaction. This was used for panels A and B. We performed serial dilution into nuclease-free water for panel C.

### Method for Fig. 3

**Fig. 3.**
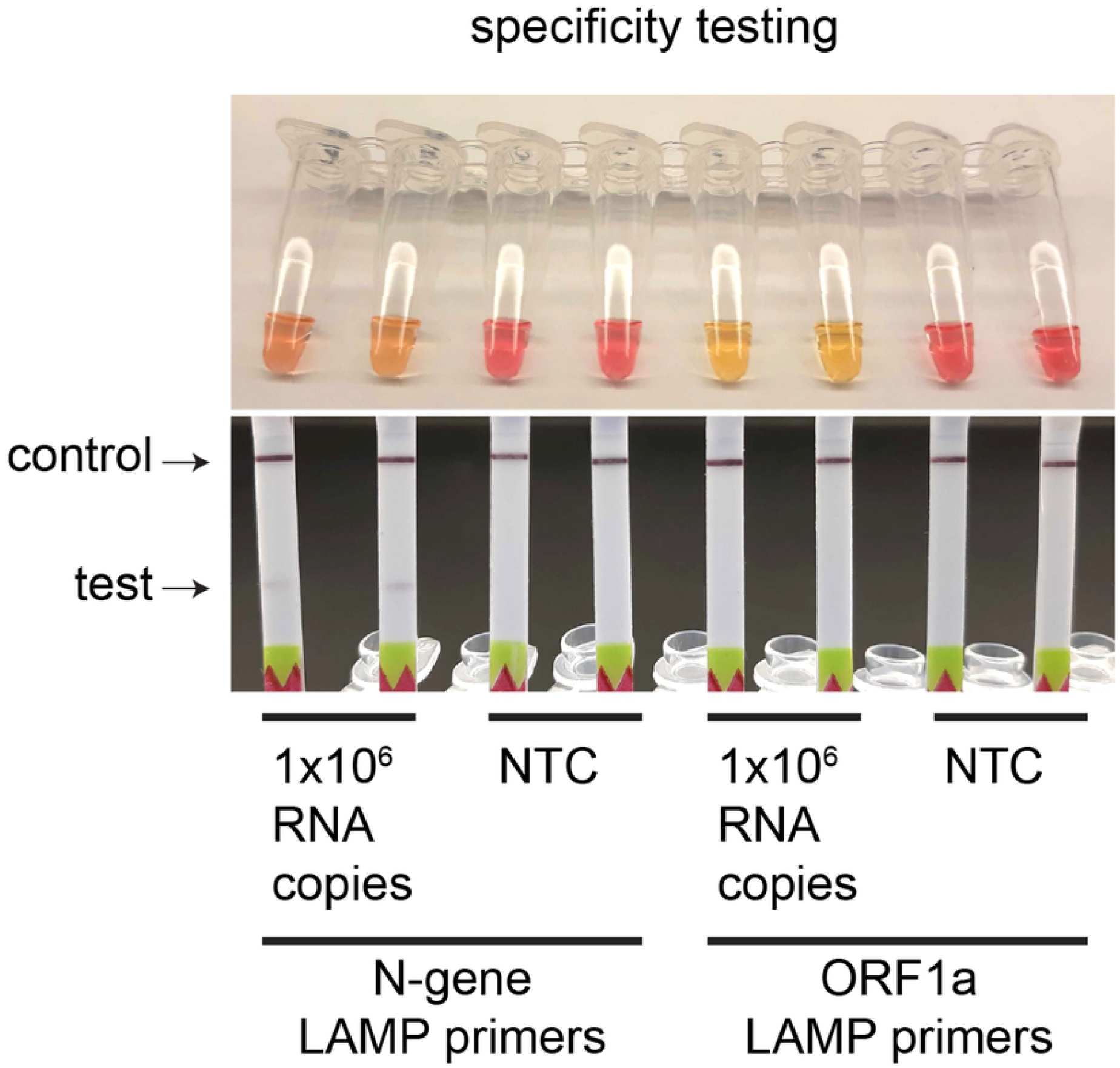
Specificity testing. Comparison between amplification run with N-gene LAMP primers (from Broughton et al., 2020, which match the ‘toehold’ of the strand-displacement probe) and ORF1a primers from Rabe and Cepko 2020 (primer set As1e, negative control, not matching the ‘toehold’ region of the strand-displacement probe). Colorimetric readout (above) shows that both primers effectively amplified the templates. Lateral flow assays (below) show no readout for ORF1a primers while the N-gene primers show positive readout as expected.

Colorimetric LAMP mix (M1800S) was used for either N-gene [ref needed] or As1e primer [ref needed] LAMP. For template, Zymo purified RNA was prepared as described for Fig. 2, diluted to 1.33×10^5^ genome/μL. Using 7.5μL of this material in a LAMP reaction gives 1 × 10^6^ viral genomes in a reaction.

### Method for Fig. 4

**Fig. 4.**
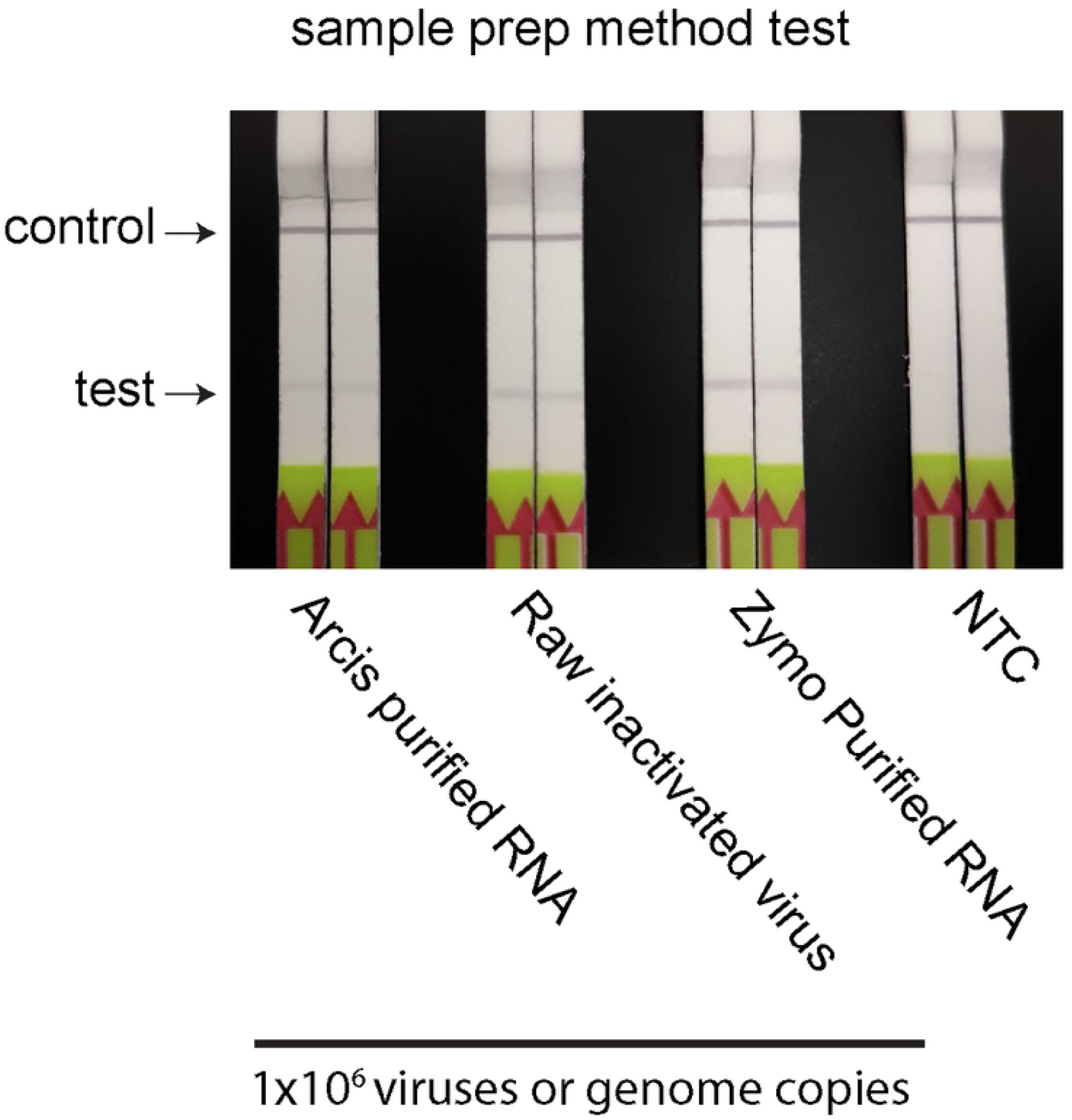
Comparison between three different forms of sample preparation. Sample preparations tested are Arcis, Inc. purified RNA; raw inactivated SARS-CoV-2 virus; and Zymo purified RNA, each at a concentration of 1×10^6^ viral genomes. NTC = No template control.

We prepared another extraction using the Arcis Coronavirus RNA Extraction Kit (Cat #UVL101) and extracted 90ul (158 ×10^6^ genomes) into a final volume of 288 μL; we then diluted this material to 1.33×10^5^ genome/μL. Using 7.5μL of this material in a LAMP reaction gives 1 × 10^6^ viral genomes in a reaction.

For raw inactivated virus we diluted the BEI Resources gamma-irradiated virus (Cat # NR-52287) 13-fold in nuclease-free water to obtain a concentration of 1.33×10^5^ genome/μL. Using 7.5μL of this material in a LAMP reaction gives 1 × 10^6^ viral genomes in a reaction.

Zymo purified RNA was prepared as described for Fig. 2, diluted to 1.33×10^5^ genome/μL. Using 7.5μL of this material in a LAMP reaction gives 1 × 10^6^ viral genomes in a reaction.

### Method for Fig. 5

**Fig. 5.**
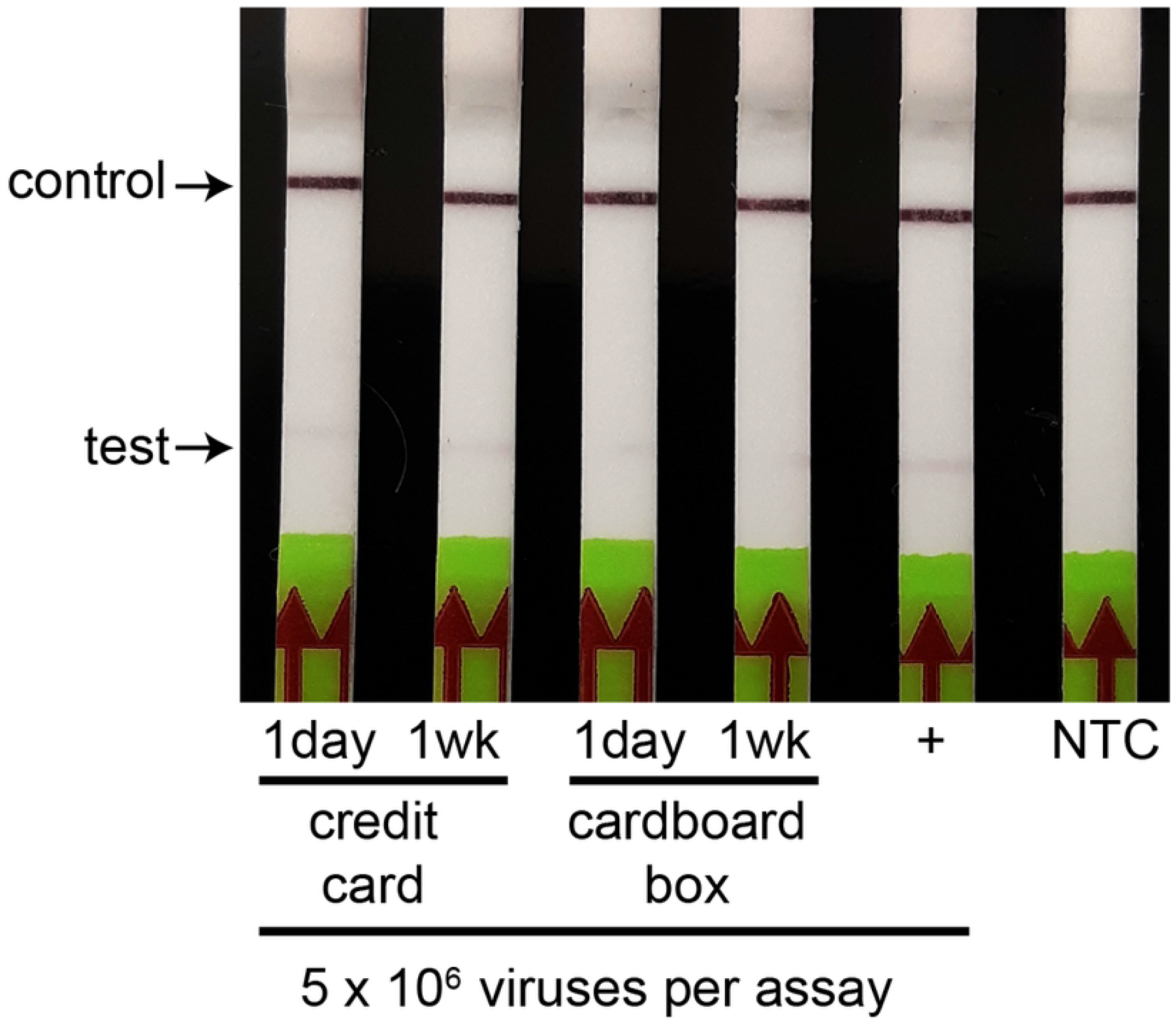
Surface testing for credit card and cardboard box after 1 day and 1 week (wk) of dry storage. A total of 5 × 10^6^ viral particles were deposited on the surfaces and dried to completion in the laminar flow hood. Either 1 day or 1 week later these dried samples were lifted off with a sterile cotton swab and PBS to perform LAMP. For positive (+) control LAMP was performed in a tube (no application to surfaces) with 5 × 10^6^ viruses. NTC = No template control.

For each surface test, 5 million inactivated viruses were spotted (in 7.5 μL) onto a plastic credit card or cardboard shipping box. The spots were allowed to dry completely for 35 minutes in a laminar flow hood. Once dry, the spots were tested on the following day or a week later (spots were circled with a pen). The dried samples were harvested from the surface by adding 200 μL PBS onto the marked area and allowed to rehydrate for 30 seconds. A sterile cotton swab was used to collect the PBS and add it to a 1.5ml eppendorf tube pre-loaded with 100 μL PBS; the cotton swab always retains PBS but this was interchanged with the PBS in the tube to yield approximately 100 μL of eluate. This eluate was used in the lamp mix (7.5 μL as outlined for Fig. 1).

## Results

We designed a strand-displacement assay for SARS-CoV-2 targeting the Broughton et al, 2020 [18] primer set (Table 1). To evaluate whether a strand-displacement probe (Fig. 1) could be used to detect SARS-CoV-2, we tested 40×10^6^ viral RNA molecules in a 30-minute LAMP reaction followed by lateral flow readout, which was successful (Fig. 2A). To examine the timeline of the LAMP amplification a fluorescence FAM readout was conducted using Zymo purified RNA at 40×10^6^ viral genomes in the reaction (Fig. 2B). The FAM readout showed a rise beginning at 15 minutes (15.8 minutes defining the threshold line) peaking at approximately 21 minutes. This suggests that the reaction time could be shortened. We hypothesize that the drop in FAM signal is tied to a lower pH as amplification products accumulate, which quenches FAM fluorescence [19] (Fig. 2B). To evaluate the sensitivity of LAMP we performed nine serial dilutions spanning from 40×10^6^ viral genomes down to 0.1 genomes per reaction (Fig. 2C). We found reliable detection down to 100 genome copies in the reaction (Fig. 2C).

To test the specificity of our lateral flow SD-LAMP assay, we tested two different LAMP primer sets. First, the As1e set (targeting ORF1a [20]) which lacks complementarity to the single-stranded ‘toehold’ [16] of the SD probe and thus serves as a negative control. Second, we used (as in Fig. 2) the Broughton et al. primer set [18] which produces a LAMP product with complementarity to the SD probe’s ‘toehold’ sequence [16]. Colorimetric readouts showed amplification of DNA using both primer sets (indicated by yellow color) while the lateral flow assays only showed positive readouts for the material amplified using Broughton et al.[18] primers (Fig. 3).

We then tested the ability of SD-LAMP to detect the virus in absence of nucleic acid extraction. The tested sample preparations were Zymo purified RNA, Arcis purified RNA and raw inactivated virus (non-purified) each at a concentration of 1×10^6^ genome copies (Fig. 4). All the tests showed positive readouts on lateral flow assays, showing that purification of RNA is not necessary to detect SARS-CoV-2, and that both RNA purification methods are compatible with the SD-LAMP assay (Fig. 4).

Finally, we tested whether we could detect dessicated SARS-CoV-2 that had been deposited onto two types of surface: cardboard (a standard shipping container) and a plastic credit card. In this experiment we deposited 5 × 10^6^ gamma-inactivated viruses onto the surfaces (7.5 μL) and allowed the spots to dry completely. After 24 hours or one week, we harvested the spots with 200 μL of PBS and a cotton swab, and then performed LAMP assays on this collected material. Remarkably, we found that these samples remain detectable, both same day and after one week, albeit with cardboard yielding consistently weaker signals (Fig. 5).

## Discussion

In this study we develop a strand-displacement probe [16,21] for the N-gene LAMP assay developed by Broughton et al [18], linking LAMP detection of SARS-CoV-2 to a lateral flow strip (Fig. 2A). We demonstrate that the method is highly specific to the Broughton et al. N-gene primers, failing to yield signal when a different primer set was used (AS1e, targeting ORF1a [20], Fig. 3) and with an LOD of 100 molecules (Fig. 2C). We also demonstrate that our method successfully detects the virus genome after 24 hours and 1 week from both non-porous (plastic credit card) and porous (cardboard box) surfaces (Fig. 5). We also show, consistent with many other studies, that RNA extraction prior to detection is not required, as we detect raw virus, along with prepared RNA from Zymo and Arcis (Fig. 4).

Our work expands the number of validated strand-displacement probes for LAMP assays, and is the first such probe for the highly popular Broughton et al.[18] primer set. While previous work on strand-displacement LAMP and SARS-CoV-2 has focused on clinical saliva samples [21], we demonstrate the utility of the method for surface detection of viral genomes, presenting new possibilities for surveillance of viral presence in the environment. In particular, the implementation of an integrated surface SARS-CoV-2 detector as a microfluidic chip would provide a cheap method to monitor viral presence on high-touch surfaces. This method is attractive as an early-warning monitor when cohorts of humans can be tested anonymously (such as high-touch surfaces in shared apartment buildings or dorm rooms, currency, or wastewater). Given that an infected individual may harbor 10^9^ viral particles per mL in their airway [22], it is reasonable to estimate that one infected person could deposit 5 ×10^6^ particles on a surface (as in Fig. 5) in only a few microliters of saliva.

Surfaces have been shown to harbor SARS-CoV-2 for hours to days; importantly, however, they are not believed to be drivers in ongoing pandemic spread. Instead, these surfaces provide valuable information about infection in a population or environment and provide a method for monitoring the movement and spread of the disease, independently of traditional disease diagnosis in individuals. Previous studies have shown that humidity and temperature modulate viral survival on surfaces [23–26] and that the virus can survive for 1-4 weeks on non-porous surfaces such as steel, glass, or plastic [26,27]. In contrast, these studies consistently suggest that viral survival on porous surfaces is lower, which has been attributed to capillary-action driven droplet spreading [28]. For example, one early study showed that SARS-CoV-2 becomes undetectable after 24h on cardboard [29]. While plastic provided more robust detection, we observed signal from cardboard surface testing even after 1 week (Fig. 5). This is surprising and may point to a hitherto underappreciated utility for even porous material as useful sentinel surfaces, and also to our innovative method of sample rehydration and recovery in PBS. Indeed, a recent LAMP study showed that plastic money cards on the Brigham Young University campus exhibited detectable virus, but its deposition and latency time was not measured as they were naturally deposited samples [30]. In light of our findings, we suggest that LAMP provides valuable surveillance for much longer than previously appreciated. Given our findings with cardboard, even paper money [31] might have more utility than previously appreciated as an early-warning system for surveillance of SARS-CoV-2.

## Conclusions

In this study we show that surface detection of SARS-CoV-2 can be performed with strand-displacement lateral-flow LAMP, and that this method retains the high sensitivity of this nucleic acid amplification method. Remarkably, we observe that surfaces can retain detectable levels of virus for up to a week, including both cardboard and plastic (porous and nonporous) surface types. Future work could include integrating the entire LAMP plus lateral flow output into a microfluidic chip design for use at surveillance sites. An additional future direction would be to leverage the high sequence descrimination power of strand-displacement probes [16] to create specific LAMP assays for the SARS-CoV-2 variants [32,33]. Since most sequence changes tend to occur in the Spike gene and other highly immunogenic regions, LAMP assays to the Nucleocapsid (N) gene (such as Broughton, et al.’s primers [18] also used in our study) can detect most SARS-CoV-2 strains and serve as pan-detectors. We note that some variants do have changes to N-gene and other regions of the genome [32,33];. nevertheless, a comprehensive, well validated suite of such assays would provide an early warning system for new variants which could then be confirmed by sequencing. In the future, LAMP and strand-displacement probes should play key roles in molecular pandemic surveillance.

## Acknowledgements

The authors acknowledge the support and advice from Dr. William Bellows, Kevin Kramer, Dr. Kathryn Walters-Conte, Dr. David O’Connor, and Dr. Shelby O’Connor. Funding was provided through the Research Expense Account of Dr. Bellows and National Science Foundation grant #1735694 to J.R.B and Kathryn Walters-Conte.

